# Dysregulation of replication stress-induced ROS in transformed cell lines: a vicious circle at cancer initiation

**DOI:** 10.1101/2025.02.27.640609

**Authors:** Sandrine Ragu, Elodie Dardillac, Sylvain Caillat, Jean-Luc Ravanat, Bernard S. Lopez

## Abstract

The canonical DNA damage response (DDR) maintains genome stability, involving DNA synthesis/cell cycle arrest. However, unchallenged cells proliferate when they are continually exposed to low-level/endogenous replication stress. We previously discovered and characterized a noncanonical cellular response that is specific to nonblocking replication stress, i.e., low-level stress (LoL-DDR), in primary cells. Although this response generates replication stress-induced reactive oxygen species (RIR), it triggers a program that prevents the accumulation of premutagenic 8-oxo-guanine (8-oxoG). Primary cells control RIR production via NADPH oxidases. Increasing the severity of replication stress above a precise threshold triggers the canonical DDR, leading to cell cycle arrest and RIR suppression, resulting in a peak-shaped dose response for RIR production. Here, we show that the LoL-DDR is dysregulated in cancer cell lines, which exhibit the following differences compared with primary cells: 1-RIR are not detoxified under high-level stress conditions, resulting in a continuous increase in the dose-response curve of RIR production; 2-RIR are not produced by NADPH oxidases; and 3-replication stress favors the accumulation of the premutagenic 8-oxoG. Moreover, using an *in vitro* breast cancer progression model, we show that LoL-DDR dysregulation occurs at an early stage of cancer progression. Since, conversely, ROS trigger replication stress this establishes a “vicious circle” involving replication-stress and ROS that continuously jeopardizes genome integrity that should fuel and amplify tumorigenesis.

## INTRODUCTION

Genome is daily assaulted by various exogenous or endogenous attacks that jeopardize its integrity, leading to cell death, inflammation, aging or tumorigenesis. To cope with these stresses, cells have developed the DNA damage response (DDR) that coordinates a network of pathways allowing the faithful transmission of the genome. In response to genotoxic stress, the well-documented “canonical” DDR (cDDR) triggers cell cycle checkpoints that slow or arrest cell cycle progression (1). Deficiency in this cDDR drives genetic instability, which is frequently associated with cancer susceptibility (1–7). Notably, the cDDR has been shown to be activated in response to replication stress at the pre/early steps of senescence and tumorigenesis (2–4, 8, 9), suggesting that this type of stress might participate in oncogenesis.

In primary cells, a threshold of stress intensity must be reached to induce the cDDR (10). In the absence of sufficiently strong stress to fully induce the cDDR, which leads to the arrest of DNA replication and cell cycle progression, cells are still routinely exposed to inevitable low stresses/endogenous stresses that can potentially also threaten genome stability. Such low/endogenous stress can include replicative and oxidative stresses. Indeed, replication fork progression can be hindered by endogenous obstacles (transcription, endogenous damage, particular DNA structures, proteins tightly bound to DNA, etc.) (11–14). Additionally, reactive oxygen species (ROS), which are spontaneous byproducts of cell metabolism, can generate premutagenic bases and can alter replication dynamics (15–17). Despite chronic exposure to low-level/endogenous stresses, cells proliferate and replicate their genome. This suggests that such low stresses escape detection by the cDDR. Recently, we discovered and characterized a “noncanonical” DDR that specifically occurs in response to low-level stress in primary human cells, and we defined this response as the low-level stress DDR (LoL-DDR); this response does not lead to the arrest of DNA replication but still protects genome integrity (10).

We have shown that primary human cells adapt their response to the severity of replicative stress in two distinct phases: full activation of the cDDR requires that stress reach an intensity threshold; below this threshold, cell proliferation is not blocked, but cells produce ROS, namely, replication-induced ROS (RIR), which then activate the detoxification program, reducing the level of the premutagenic base 8-oxo-guanine (8-oxoG) in the genome (10). Because ROS can negatively affect genome integrity, RIR production is precisely controlled by cells, and RIR production follows known trends in primary human cells. 1-The dose response of RIR follows a peak-shaped curve; RIR production increases with replication stress intensity until a threshold is reached, and above this threshold (when the stress intensity increases), the cDDR is induced, cell progression is arrested, and RIR are scavenged, resulting in a peak-shaped dose‒response curve for RIR. 2-RIR production is controlled by the NADPH oxidases DUOX1 and DUOX2. 3-RIR prevents the accumulation of 8-oxoG in the genome in an adaptive manner (10). Therefore, in primary cells, RIR appear to function as secondary messengers of an autonomous response to preserve genome stability, specifically in response to low-level/endogenous stress (LoL-DDR), which do not lead to the arrest of replication progression. These findings reveal the precise regulation of primary cell responses to stress severity. This response is critical because, in contrast to severe stress conditions, cells are exposed to chronic pernicious and inevitable low-level/endogenous stress conditions each day throughout their lifespan.

Because genetic instability is a hallmark of cancer cells (7, 18, 19), we addressed the question of whether the LoL-DDR might be dysregulated in transformed cell lines. Here, we show that in immortalized/transformed cells, 1-RIR production follows a dose‒response curve, continuously increasing with stress severity, even at higher doses, which differs from with the peak-shaped dose‒response curve observed in primary fibroblasts in which RIR production decreases in response to higher stress; 2-RIR are not produced by NADPH oxidases in transformed cells, unlike in primary cells; and 3-replication stress in transformed/immortalized cells leads to the accumulation of the premutagenic lesion 8-oxoG into the genome, which differs from findings in primary fibroblasts. Moreover, in an *in vitro* cellular model, the LoL-DDR appears to be altered at an early stage of breast cancer progression.

These data reveal the dysregulation of the LoL-DDR in transformed cells, which might reveal the following potential sequence of events: replication stress induces ROS production, which results in the accumulation of premutagenic lesions in the genome; reciprocally, ROS induce replication stress (15, 17). Therefore, these data reveal a “vicious circle”. This cycle may maintain and amplify genetic instability. This vicious circle” might be critical for the initiation/progression of cancer, which is consistent with the fact that transformed cells have highly active replication programs and high levels of ROS and with previous description of replication stress *in vivo* in pre/early stages of oncogenesis (2–4, 8, 9).

## RESULTS

### The dose‒response curve of RIR production is altered in transformed cell lines

First, we measured the intracellular levels of ROS by FACS using the 2’, 7’-dichlorofluorescein diacetate (DCFDA) fluorescent probe **(Figure 1A**), which allows convenient analysis of the dose response to a treatment that induces replication stress (10). In primary fibroblasts or epithelial cells, the RIR dose‒response curve exhibited a peak shape; maximum RIR production was observed in response to 250 µM hydroxyurea (HU), which is an inhibitor of ribonucleotide reductase that results in an imbalance of the nucleotide pools and *in fine* replication stress. At higher replication stress (HU> 250 µM) a cell cycle progression and the level of RIR decreases (see (10) and the scheme in **Figure 1B**). Importantly, this type of peak-shaped curve has been described in 4 different primary fibroblast strains and in primary epithelial cells, for each of the individual repeat of the experiment, at the same doses of replication stress-inducing drugs, attesting the robustness of the response in primary human cells (10).

**Figure 1.**
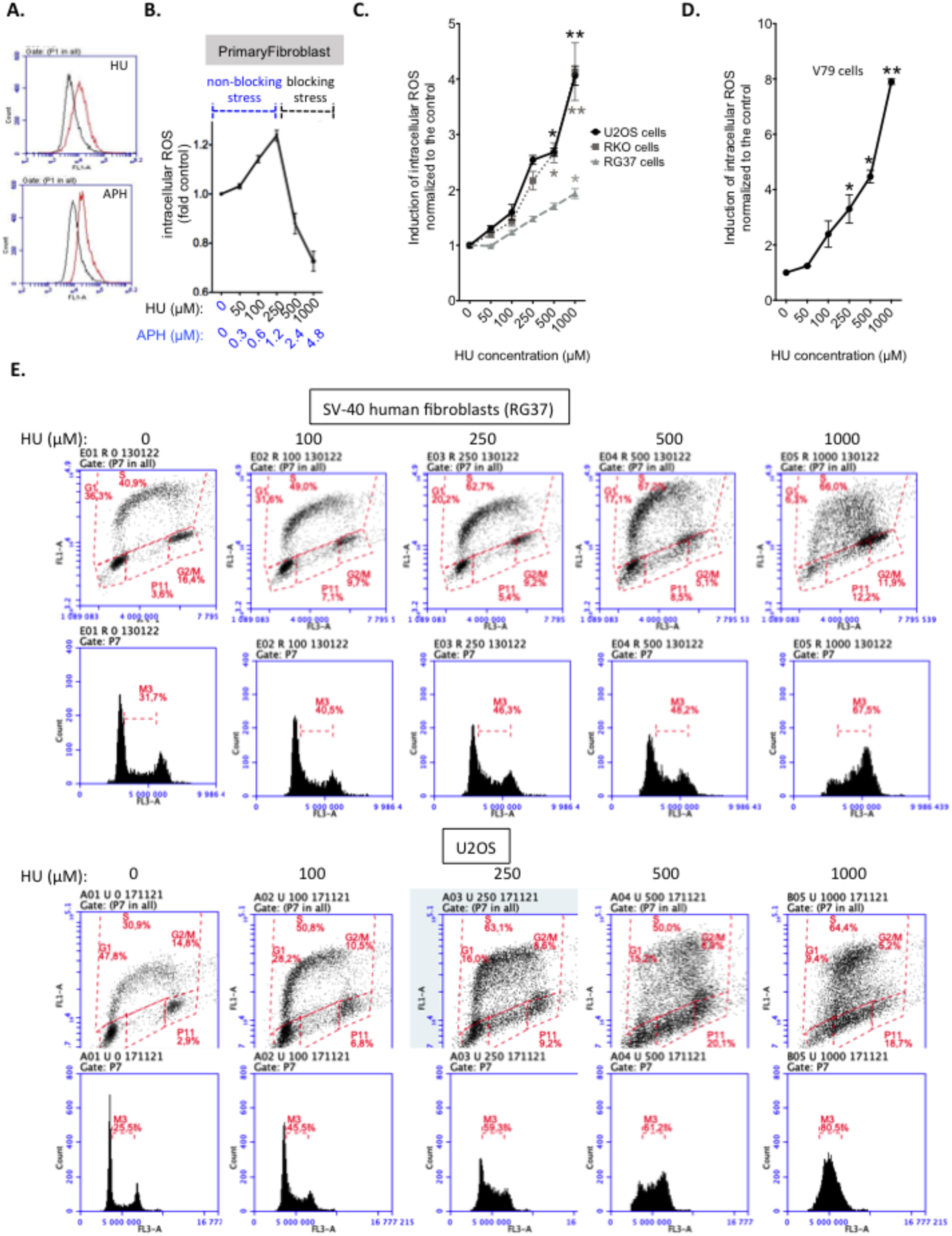
Primary and transformed cells differentially produce ROS in response to replication stress. **A**. HU- or APH-induced ROS production was monitored using the DCFDA fluorescent probe and FACS analysis. The shift in the fluorescence peak reveals the induction of intracellular ROS production. **B.** Typical peak-shaped dose-response curve of primary fibroblasts or epithelial cells following replication stress (10). **C.** Dose response of ROS induction in three different human cell lines after 48 h of exposure to HU. **D.** Dose response of ROS induction in V79 hamster cells after 48 h of exposure to HU. **E**. Impact of exposure to HU on the cell cycle distribution. The data from at least three independent experiments are presented as the mean (± SEM) level of ROS production normalized to that of the control. The non-parametric Kruskal-Wallis test was used.

In parallel with the experiments with human primary fibroblasts described above (10), we treated different transformed cell lines with increasing doses of drugs that generate replication stress. After exposure to HU, all three human cell lines, namely, the U2OS (human osteosarcoma), RKO (human colon cancer with wild-type p53), and RG37 (SV40-tranormed human fibroblasts (20)) cell lines, responded similarly; that is, these cells lines exhibited a gradual increase in RIR production, but in contrast to primary cells (10), there was no decrease in RIR product after treatment with higher doses of HU **(Figure 1C and** compared with **Figure 1B** and (10)). A hamster cell line, namely, the V79 cell line (a p53-mutated lung Chinese hamster cancer cell line), also exhibited the same behavior (**Figure 1D**). This finding shows that the same dose-dependent continuous increase in RIR production is observed in several types of cancer cell lines, even those with different p53 statuses and those from different species (human and hamster). These curves are different from those observed in primary cells (compare **Figure 1B to Figure 1C–D** and (10)). In primary fibroblasts, RIR are abolished at high HU doses (500 or 1000 µM) when DNA replication is blocked (10). In transformed cells, RIR production still increased at such high doses (**Figure 1C-D**), i.e., even when replication was strongly abrogated and when the cells accumulated in the S or G2/M phase (**Figure 1E**). RIR production thus appears to be dysregulated in transformed cells compared with primary cells, reflecting a change in regulation of the Lol-DDR. Therefore, the dose‒response curves of RIR production represent a reliable diagnosis marker of changes in the LoL-DDR. Using this readout, we then analyzed other parameters.

We next investigated another replication stress inducer, namely, aphidicholin (APH), which is an inhibitor of replicative polymerase. Exposure of primary cells to APH also resulted in a peak-shaped response in RIR production, which peaked after treatment with APH doses between 0.6 and 1.2 µM and then decreased after treatment with higher doses (**Figure 1B** and (10)). In contrast to primary cells, but similar to HU treatment, exposure of all 3 human transformed cell lines to APH led to gradual continuous RIR production, with increased ROS production observed even after treatment with higher doses (**Figure 2A**). Together, these data suggest that RIR product actually occurs due to replication stress but that transformed cell lines respond differently than primary cells.

**Figure 2.**
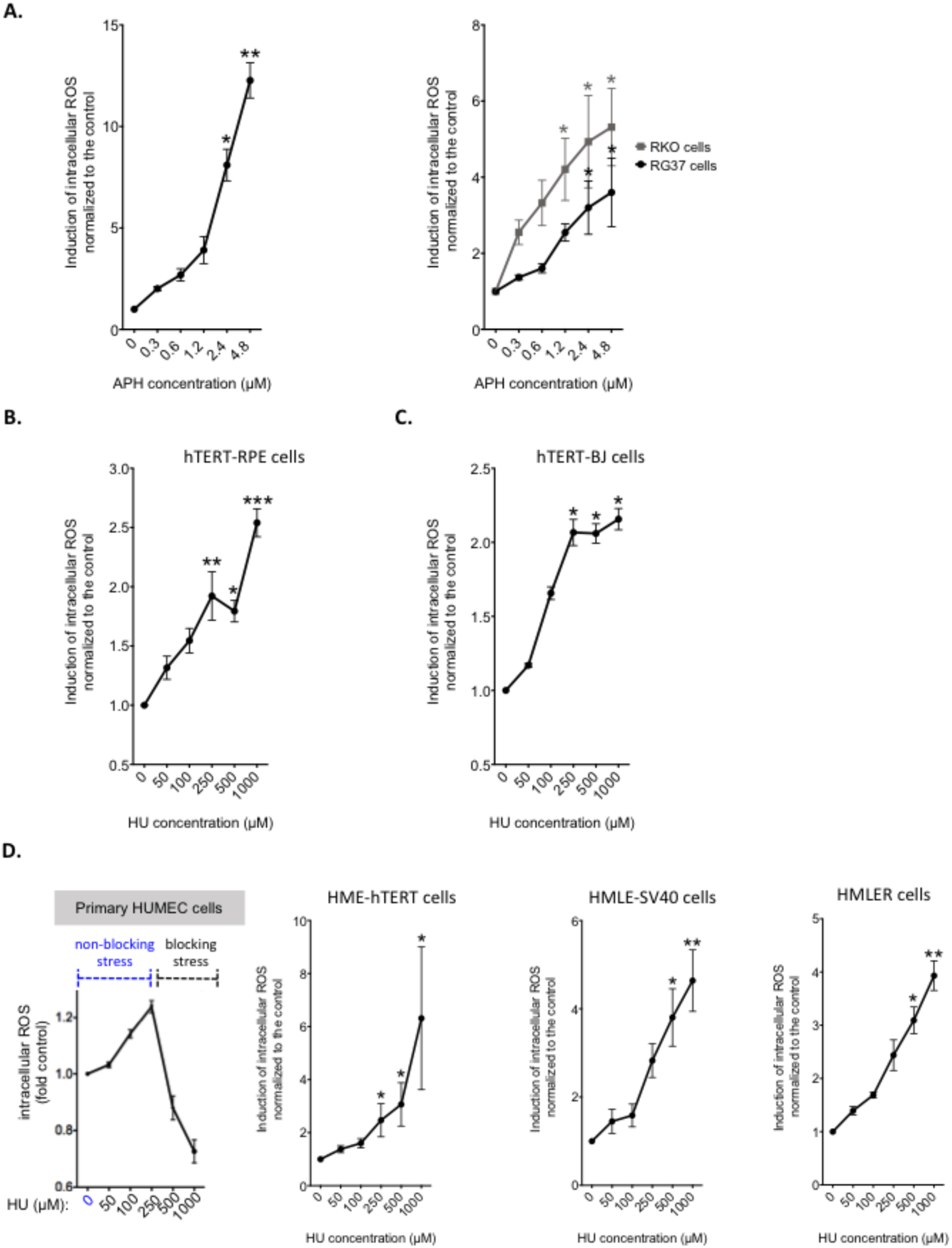
RIR production under different conditions. **A.** Impact of APH exposure on three different human transformed cell lines. **B.** RIR production in response to HU in RPE-hTert immortalized cells. **C.** RIR production in response to HU in human fibroblast-hTert immortalized cells. **D.** RIR production in response to HU in the *in vitro* breast cancer progression model. The data from at least three independent experiments are presented as the mean (± SEM) level of ROS production normalized to that of the control. The non-parametric Kruskal-Wallis test was used.

Telomerase-immortalized retinal pigment epithelium (RPE) cells exhibited a dose‒ response curve similar to that of the cell lines described above, i.e., a continuous and progressive increase in RIR production was observed, although the level of stimulation was lower (**Figure 2B**). However, human fibroblasts that were immortalized with telomerase (BJ-hTert) exhibited a response that was intermediated relative to the responses of primary and transformed cells (**Figure 2C**). Indeed, after treatment with higher HU doses, the RIR levels neither increased nor decreased; rather, RIR levels plateaued (**Figure 2C**). This finding shows that while the LoL-DDR is actually altered in immortalized cells, such alteration differs in cells of different tissue types. Nevertheless, these data suggest that changes in the LoL-DRR might occur in the early stage of cell transformation.

### Dysregulation of the LoL-DDR occurs at an early stage of cancer progression in an *in vitro* cell model

To address the above question, we used a cell culture model of breast cancer progression (21, 22). This model was established with primary epithelial cells (HUMECs), telomerase-immortalized HUMECs (HME-hTert cells), HME-hTert cells expressing the SV40 large T antigen (HMLE-SV40), and HMLE-SV40 cells expressing the V-RAS oncogene. Notably, RIR levels peaked in primary breast epithelial cells after treatment with 250 µM HU (**Figure 2D** and (10)), which is similar to what was observed in primary fibroblasts (see **Figure 1B** and (10)). The dose‒response curves of the immortalized HME-hTert cells lost their peak shape, and RIR production progressively increased with increasing dose (**Figure 2D**), similar to what was observed in the other transformed cell lines (compare with **Figure 1C-D**). Notably, the change in dose response occurred as soon as the HMR-hTert cells, and the curve increased with increasing dose. This response is similar to that of RPE-hTert cells (compare with **Figure 2B**), another epithelial cell line. These data suggest the potential dysregulation of the LoL-DDR in an early stage of breast cancer progression.

### RIR are not produced by NADPH oxidases in immortalized/transformed cell lines

In primary fibroblasts, RIR are secondary messengers of the LoL-DDR, and they protect genome stability in an adaptive way (10). As such, RIR production is precisely controlled by the cellular enzymes NADPH oxidases (10). The exposure of primary human cells (fibroblasts as well as epithelial mammary cells) to diphenylene iodonium chloride (DPI), which inhibits all NADPH oxidases (23), suppressed RIR production (10). In contrast to its effect on primary cells, DPI did not reduce RIR levels in transformed cells, and it even show a trend to induce ROS production in cells that were not exposed to HU (**Figure 3A**). Moreover, in the breast cancer progression model used here, although RIR production was indeed abrogated by DPI in primary epithelial cells (HUMEC), RIR production became resistant to DPI as early as the HME-hTert step (**Figure 3B**), i.e., in the same stage as when the alteration in the dose‒response curves of RIR production was observed. Collectively, these data confirm the dysregulation of the LoL-DDR in an early stage of breast cancer progression, at least in *in vitro* cellular models.

**Figure 3.**
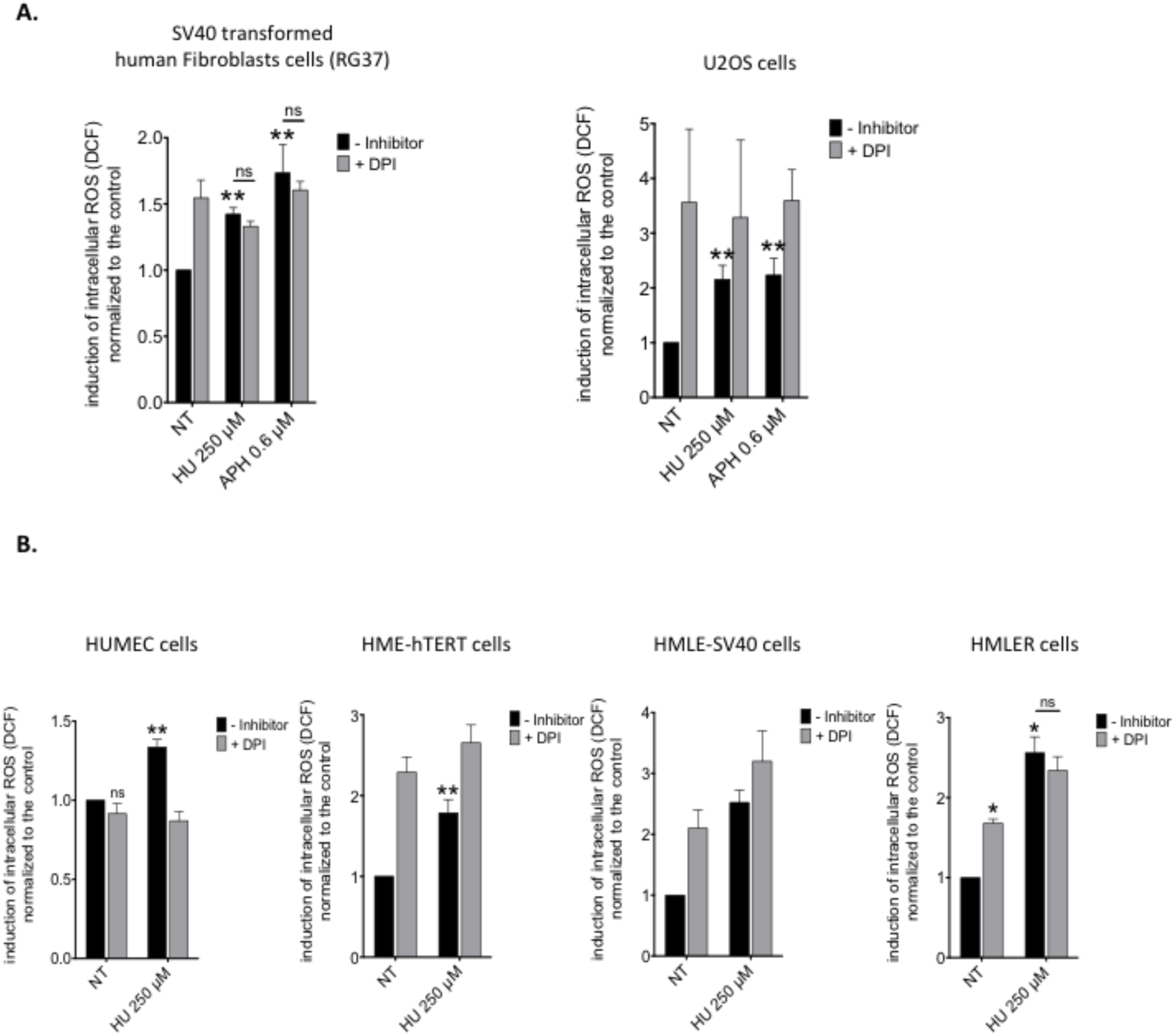
RIR are resistant to NADPH oxidase inhibition in immortalized/transformed cells. **A.** U2OS and SV40-transformed human fibroblasts**. B.** *In vitro* breast cancer progression model. The data from at least three independent experiments are presented as the mean (± SEM) level of ROS production normalized to that of the control. **C.** Telomerase-immobilized human fibroblasts (BJ-hTert).

The total cellular levels of RIR were not affected by DPI, which suggests that no or only a marginal portion of RIR might be produced by NADPH oxidases and that ROS from other cellular sources account for a large portion of the cellular RIR levels in transformed cell lines.

### Replication stress leads to the accumulation of oxidative DNA lesions in transformed cell lines

In primary cells, RIR prevent the accumulation of oxidative DNA damage in the nuclear genome (10). Given the alteration in the LoL-DDR that was observed in transformed cells, we assessed whether RIR might differentially impact the accumulation of oxidative damage in the nuclear genome. To address this question, levels of the main premutagenic oxidized base lesion, namely, 8-oxoG, which is a marker of genotoxic oxidative stress, were quantified through isotope dilution high-performance liquid chromatography coupled with electrospray ionization tandem mass spectrometry (HPLC–MS/MS) (24). In primary cells, the frequency of 8-oxoG decreased at doses that induced peak RIR production (50 and 250 µM HU), and then, 8-oxoG did not further accumulate after treatment with higher HU doses (**Figure 4A**) (10). In contrast, in transformed cells, the frequency of 8-oxoG in genomic DNA gradually increased upon HU treatment in a dose-dependent manner (**Figure 4B**), which was consistent with the dose‒ response curves of RIR production (compare **Figure 4B** with **Figure 1C-D**). Consistently, in TERT-immortalized fibroblasts (BJ-Tert), the level of 8-oxoG increased and plateaued after treatment with the HU dose that caused the RIR levels to plateau (compare **Figure 4C with Figure 2C**).

**Figure 4.**
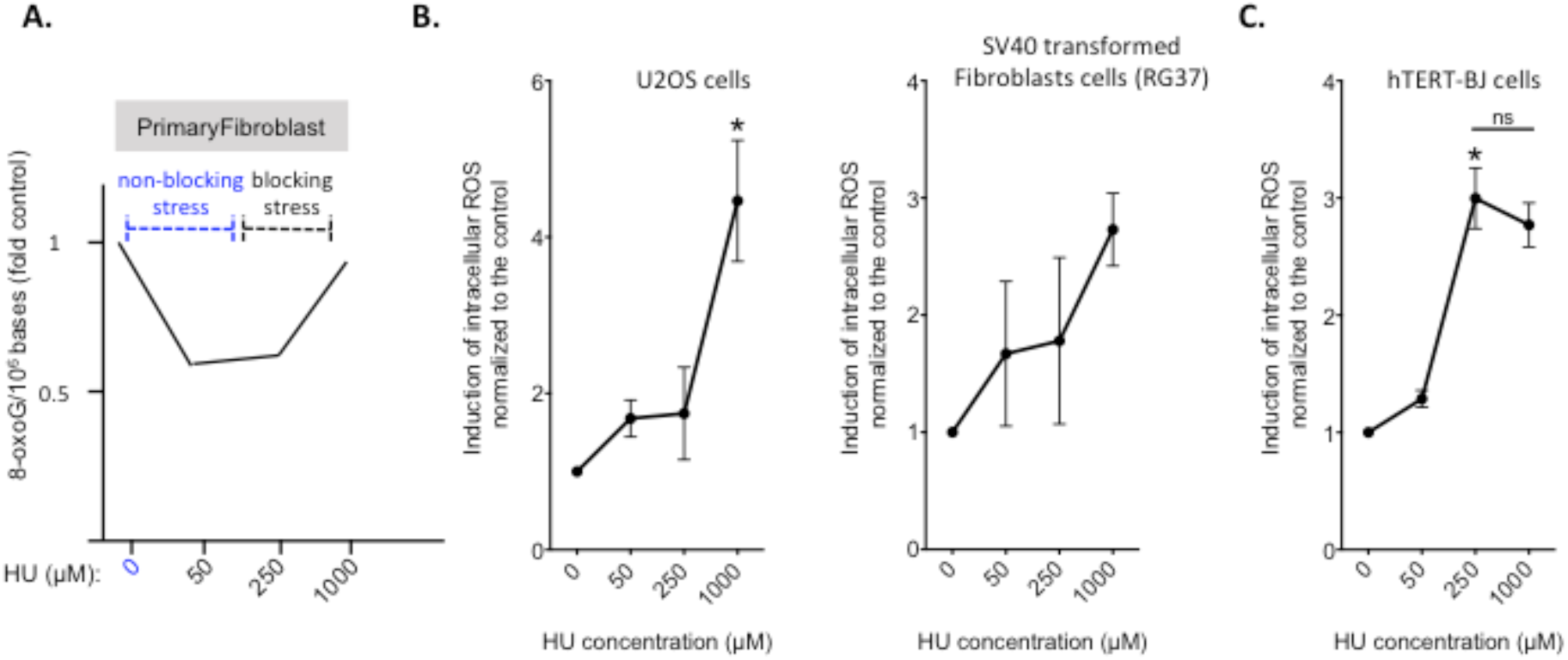
8-oxoG accumulation. **A.** Primary fibroblasts, data from (10). **B**. U2OS- and SV40-transformed human fibroblasts. **C**. human telomerase-immortalized fibroblasts BJ-Tert. The data from three independent experiments are presented as the mean (± SEM) level of ROS production normalized to that of the control. The non-parametric Kruskal-Wallis test was used.

Collectively, these data support the concept that the LoL-DDR, which prevents the accumulation of premutagenic lesions in primary cells in response to low-level/endogenous replication stress (10), is altered in transformed cells.

## DISCUSSION

Cells are routinely exposed to stresses that compromise the integrity of their genome. The cDDR allows cells to cope with these genotoxic stresses. Schematically, the cDDR stops cell cycle progression, providing time and cofactors access to the repair machineries. This enables the resumption of the cell cycle with restored DNA matrices (25–27). Defects in the cDDR lead to genome instability and cancer predisposition (2, 4–7, 25). However, in the absence of strong exogenous stresses, cells can divide and replicate their genome despite the fact that they are faced with endogenous stress conditions. Recently, we showed that a threshold of stress intensity is required to induce the cDDR in primary human cells (10). We identified and characterized a noncanonical DDR specific to low-level stress (LoL-DDR) in primary human cells, which results in RIR generation but prevents the accumulation of the premutagenic lesion 8-oxoG in the genome in an adaptive manner (10). This pathway is characterized by the fact that: 1-the level of RIR decreases at high level stress intensity, 2-RIR are produced by the NADPH-oxidases, and 3-the prevention against the accumulation of pre-mutagenic oxidative lesions such as 8-oxoG (10). Here, we show that the protective LoL-DDR is dysregulated in transformed cell lines. 1-RIR still accumulate under high replication stress conditions, which differs from RIR levels in primary cells; 2-cellular RIR production occurs independently of NADPH oxidase control; and 3-the accumulation of premutagenic 8-oxoG in the cell genome. In addition, experiments using an *in vitro* cellular model of breast cancer progression suggested that dysregulation of the LoL-DDR occurs in an early stage of cancer progression.

An increase in premutagenic lesions with increased replication stress should induce mutagenesis, resulting in genetic instability that could foster cancer initiation and progression as well as the appearance of neoantigens. Importantly, the process described here has been shown to occur in cell lines with wild-type p53 (RKO, U2OS, and RPE-tert cells) as well as in cells with mutated p53 (V79 cells) or in which p53 is trapped by the SV40 large T antigen (RG37 cells). Thus RIR production in immortalized/transformed cell lines occurs independently of p53. These data are consistent with the fact that LoL-DDR dysregulation occurs in an early stage of cancer progression, as shown here.

As summarized in **Figure** 5, in primary noncancer cells, nonblocking, low-level replication stress induces the LoL-DDR, which protects genome stability; increasing the stress intensity turns off the LoL-DDR but induces the cDDR, which maintains genetic stability but in a different way. Notably, all the factors that play pivotal roles in controlling the LoL-DDR, are lost in different types of tumors (28–34). In immortalized cells, the LoL-DDR is dysregulated in the early stage of cancer progression, and replication stress induces ROS production even at high levels of stress, leading to the accumulation of premutagenic lesions in the genome. Since, ROS cause replication stress (15, 17), this triggers a vicious cycle, which maintains and amplifies genetic instability and thus should play an important role in cancer etiology in the both initiation and progression stages (**Figure 5**).

**Figure 5.**
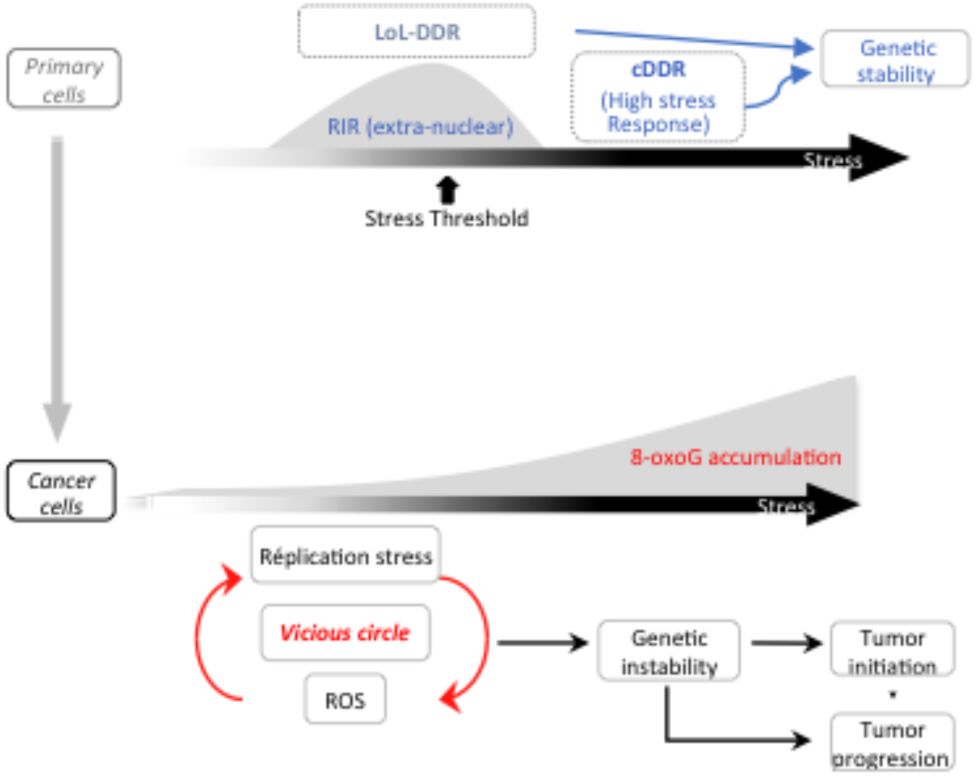
Different responses of cancer and primary cells to replication stress. Upper panel: Below a particular threshold of stress intensity, primary cells induce the LoL-DDR, which is a noncanonical stress response that produces RIR, triggering detoxification in an adaptive way. Above this threshold, the cells induce the cDDR and detoxify RIR (10). Both responses protect genome stability. Lower panel: In cancer cells, these responses are dysregulated, and ROS levels continuously increase with increasing stress intensity, resulting in the accumulation of premutagenic oxidative lesions (8-oxoG). Since ROS induce replication stress (15, 17), this reciprocally triggers vicious circle of replication stress and ROS production, which promote genome instability, potentially fueling tumor initiation and progression.

## MATERIALS AND METHODS

### Cell culture and treatments

The cells were grown at 37 °C with 5% CO_2_ in DMEM supplemented with 10% fetal calf serum (FCS).

Primary human mammary epithelial cells (HuMECs) were obtained from Thermo Fisher Scientific and cultured according to the manufacturer’s recommendations. HuMEC derivatives (HME, HMLE, and HMLER cells) were obtained from Professors Alain Puisieux (Lyon, France) and Robert Weinberg (Boston, USA) and were cultured in 1:1 DMEM/HAM-F12 medium (Gibco, Life Technologies) supplemented with 10% FBS (Lonza Group, Ltd.), 100 U/ml penicillin‒streptomycin (Invitrogen), 10 ng/ml human epidermal growth factor (EGF) (Sigma), 0.5 µg/ml hydrocortisone (Sigma) and 10 µg/ml insulin (Sigma). Primary fibroblasts were exposed to HU, APH or CPT for 72 h at 37 °C. For antioxidant treatment, primary fibroblasts were exposed to 2 mM NAC (Sigma‒Aldrich, St. Louis, MO, USA) for 72 h.

### Measurement of cellular ROS production by FACS analysis

Cellular ROS production was measured with a CM-H2DCFDA (2’, 7’-dichlorofluorescein diacetate) (Life Technologies, USA) or dihydrorhodamine 123 (DHR 123) (Sigma) assay kit according to the manufacturer’s protocol. Approximately 10^5^ cells/well were plated into 6-well plates and incubated at 37 °C in 5% CO_2_. After 2 days, the cells were washed with PBS and incubated with 10 µM CM-H2DCFDA or DHR 123 in DMEM supplemented with 1% FBS for 45 min at 37 °C in the dark. The cells were trypsinized and resuspended in DMEM supplemented with 1% FBS. The pelleted cells were washed again, and the live cells were resuspended in PBS and analyzed on a BD Accuri C6 flow cytometer (BD Biosciences, San Diego, CA) equipped with an FL1 laser (515–545 nm). The data are presented as the mean percentages of four independent experiments.

### Cell cycle analysis and BrdU incorporation

Cells were incubated in culture medium supplemented with or without HU for 72 h at 37 °C, and 5-bromo-2-deoxyuridine (BrdU, Sigma) was added to the culture medium at a final concentration of 10 µM and incubated for 30 min. Pelleted cells were detached with trypsin, fixed with 80% ethanol, and resuspended in 30 mM HCl/0.5 mg/ml pepsin. BrdU was immunofluorescently labeled with a mouse anti-BrdU antibody (DAKO, clone Bu20a) and a fluorescein-conjugated donkey anti-mouse antibody (Life Technologies), and the cells were stained with propidium iodide (PI; 25 µg/ml) in the presence of ribonuclease A (50 µg/ml). Flow cytometry analyses were performed using an Accuri C6 flow cytometer (BD Biosciences).

### 8-OxoG measurement

Genomic DNA was extracted and enzymatically digested using an optimized protocol that minimizes DNA oxidation during the procedure (35). Then, 8-oxoG levels were quantified using isotope dilution high-performance liquid chromatography coupled with electrospray ionization tandem mass spectrometry (HPLC-MS/MS) as previously described (24); ^15^N_5_-8-oxoG was used as the internal standard. In addition to the mass spectrometric detector, the system is equipped with a UV detector that is set to 260 nm to measure the quantity of normal nucleosides. The results are expressed as the number of 8-oxoG lesions per million normal nucleosides.

### Statistical analyses

Statistical analyses were performed with GraphPad Prism 10. P < 0.05 was considered to indicate a statistically significant difference.

## ACKNOWLEDGMENTS

We thank Corinne Dupuy (Gustave Roussy Cancer Institute) for helpful discussions and advice on NADPH oxidases.

## Declaration of Interests

The authors declare that they have no conflicts of interest.

## Author Contributions

Conceptualization: BSL; funding acquisition: BSL; investigation: SR, ED, SC; methodology: SR, BSL, SC, and J-LR; 8-oxoG dosage experiments: SC and J-L R; project administration: BSL; resources: BSL; supervision: BSL; validation: SR, ED, J-LR, and BSL; writing the original draft: BSL; and writing – reviewing & editing: all the authors.

## Funding

This work was supported by grants from the Institut National du Cancer (PLBIO21-072, PLBIO24-194), La Ligue Nationale Contre Le Cancer (ARN thérapeutiques), Ligue régionale contre le cancer (comité Ile de France), (ITMO Cancer (PCSI 2022), and Fondation ARC (ARCPJA2022060005157).

## Notes

### Competing Interest Statement

The authors have declared no competing interest.

